# Selective stimulation of the ferret abdominal vagus nerve with multi-contact nerve cuff electrodes

**DOI:** 10.1101/2021.02.19.431879

**Authors:** Jonathan A. Shulgach, Dylan W. Beam, Ameya C. Nanivadekar, Derek M. Miller, Stephanie Fulton, Michael Sciullo, John Ogren, Liane Wong, Bryan McLaughlin, Bill J. Yates, Charles C. Horn, Lee E. Fisher

**Affiliations:** Dept. of Biomedical Engineering, Carnegie Mellon University, Pittsburgh, PA 15213; Dept. of Otolaryngology, University of Pittsburgh, Pittsburgh, PA 15213; Dept. of Bioengineering, University of Pittsburgh, Pittsburgh, PA 15213; Center for Neural Basis of Cognition, Pittsburgh, PA 15213; UPMC Hillman Cancer Center, University of Pittsburgh, Pittsburgh, PA 15213; Micro-Leads Inc., Somerville, MA 02144; Dept. of Neuroscience, University of Pittsburgh, Pittsburgh, PA 15213; Center for Neuroscience, University of Pittsburgh, Pittsburgh, PA 15213; Dept. of Medicine, University of Pittsburgh, Pittsburgh, PA 15213; Dept. of Anesthesiology, University of Pittsburgh, Pittsburgh, PA 15213; Dept. of Physical Medicine & Rehabilitation, University of Pittsburgh, Pittsburgh, PA 15213

**Author notes:** Authors JAS and DWB contributed equally to this work. To whom correspondence should be addressed: Lee E. Fisher, 3520 Fifth Avenue, Suite 300, Pittsburgh, PA 15213.

## Abstract

Dysfunction and diseases of the gastrointestinal (GI) tract are a major driver of medical care. The vagus nerve innervates and controls multiple organs of the GI tract and vagus nerve stimulation (VNS) could provide a means for affecting GI function and treating disease. However, the vagus nerve also innervates many other organs throughout the body, and off-target effects of VNS could cause major side effects such as changes in blood pressure. In this study, we aimed to achieve selective stimulation of populations of vagal afferents using a multi-contact cuff electrode wrapped around the abdominal trunks of the vagus nerve. Four-contact nerve cuff electrodes were implanted around the dorsal (N=3) or ventral (N=3) abdominal vagus nerve in six ferrets, and the response to stimulation was measured via a 32-channel microelectrode array (MEA) inserted into the nodose ganglion. Selectivity was characterized by the ability to evoke responses in MEA channels through one bipolar pair of cuff contacts but not through the other bipolar pair. We demonstrated that is was possible to selectively activate subpopulations of vagal afferents using abdominal VNS. Additionally, we quantified the conduction velocity of evoked responses to determine what types of nerve fibers (i.e. Aδ vs. C) responded to stimulation. We also quantified the spatial organization of evoked responses in the nodose MEA to determine if there is somatotopic organization of the neurons in that ganglion. Finally, we demonstrated in a separate set of three ferrets that stimulation of the abdominal vagus via a four-contact cuff could selectively alter gastric myoelectric activity, suggesting that abdominal VNS can potentially be used to control GI function.

## Introduction

The abdominal vagus nerve is involved in the control of gastrointestinal (GI) tract motility and inflammation, as well as pancreatic function, and consequently plays critical roles in GI disease, obesity, and diabetes^1^. The mechanisms through which vagal pathways mediate these effects, however, have been difficult to determine due to complex organ innervation, and also because the abdominal vagus nerve contains a variety of fiber types, of which approximately 80% are afferent fibers and 20% are efferent fibers^2^. Strategies for understanding vagal function have relied primarily on pharmaceutical and ablation approaches but with limited success^3,4^. Vagus nerve stimulation (VNS) with electrical current has been used to interrogate vagal function, as well as for therapeutic applications, including control of epileptic seizures and obesity^5,6^. However, the therapeutic effects of existing VNS approaches have been modest, with significant off-target effects, including hoarseness, cough, dyspnea, pain, paresthesia, nausea, and headache^7^. Importantly, available clinical VNS systems use electrodes with contacts that fully wrap around the nerve and simultaneously engage many functional pathways, thus providing little insight into mechanisms of action of nerve stimulation and limiting the potential to tune VNS to limit side-effects^8^.

Many studies have demonstrated that multi-contact nerve cuff electrodes, with contacts spaced around the circumference of the nerve, can achieve a selective interface with peripheral nerves, allowing for targeting of specific functions while avoiding off-target effects^9^. For example, studies have shown that selective stimulation of the median, radial, and ulnar nerves can activate particular muscles of the hand and arm^10,11^. Often, selectivity of stimulation of efferent neurons is measured by recording evoked EMG activity in skeletal muscles^11,12^ or by measuring joint torques in response to muscle contractions^13,14^. For studies focused on stimulation of sensory afferents, selectivity can be measured by psychophysical measurement of perception of those sensations^15^, EMG recording of reflexive muscle responses to stimulation^16^, or by recording compound action potentials via nerve cuff recordings^17^. Measuring the selectivity of VNS is challenging because the physiological responses to stimulation are often not well understood or may only occur following minutes or hours of stimulation; for example, changes in immune responses or gastrointestinal rhythms^18^. Techniques that quantify selectivity by measuring effects on end-organ function or by recording evoked responses in nerve branches are also highly challenging because the vagus nerve branches are small, variable, and often inaccessible; they also innervate organs throughout the abdomen^19^.

The nodose ganglion is a compelling location to measure the selectivity of VNS, particularly for stimulation of the abdominal vagus nerve. The nodose ganglia are enlargements of the vagus nerve containing the cell bodies of autonomic sensory neurons projecting from organs throughout the body including the heart, lungs, and alimentary tract^20,21^. Action potentials can be recorded from these cell bodies using microelectrode arrays (MEA)^22^.

In this study, we sought to assess if specific populations of axons in the abdominal vagus nerve could be selectively activated using VNS. Experiments were conducted in ferrets, which have several advantages as a model for assessing abdominal vagal function because of similarity to humans in gastric anatomy, vagal regulation of gastric motility, and emetic responses^23^. In six animals, a four-contact cuff electrode was wrapped around the abdominal vagus nerve and compound action potential (CAP) signals elicited by stimulation through different contact pairs were recorded using a 32-channel MEA inserted in the nodose ganglion. We varied stimulation parameters (pulse amplitude and pulse width) to ascertain the maximal number of MEA channels that responded to stimulation through one cuff electrode pair or the other. We also measured the conduction velocity of the nerve fibers that were activated by stimulation, and quantified the somatotopic organization of the responses in the nodose ganglia through a nearest-neighbor analysis. Finally, we demonstrated that our abdominal VNS approach can selectively drive changes in gastric myoelectric activity, demonstrating the promise of this technique for treating GI diseases.

## Results

### Overview

The primary goal of this study was to determine whether subpopulations of axons in the abdominal vagus nerve could be selectively activated as a means to control GI function while avoiding off-target side effects. In six ferrets, a cuff electrode with two bipolar pairs of contacts (Figure 1) was wrapped around the dorsal (N=3) or ventral (N=3) abdominal vagus nerve, and the response to stimulation was measured by recording evoked CAPs through a 32-channel MEA implanted into the nodose ganglion (left for ventral vagus nerve and right for dorsal vagus nerve, as these are the predominant pathways for neurons innervating the ventral and dorsal regions of the stomach, respectively)^24^. Stimulation amplitude (0-3000 μA) and pulse width (0.1, 0.5, and 1.0 ms) were varied using a binary search algorithm to quantify threshold (i.e., the minimum stimulation amplitude that evoked a response in the nodose for each pulse width) and their effect on selectivity of VNS. For each unique amplitude/pulse width combination, a train of 120 pulses was delivered through a bipolar cuff contact pair at 2 Hz. A stimulus-triggered average of the response recorded from each MEA channel was calculated to reduce noise and facilitate detection of CAPs (Figure 2), which often had a peak-to-peak amplitude of less than 10 μV. For each pulse width, the binary search varied stimulation amplitude values with a 20 μA target resolution. If a response was detected on any MEA channel at a stimulation amplitude, amplitude would be decreased by half the difference from the next lowest amplitude tested, and if no response was detected, amplitude would be increased by half the distance from the next highest amplitude tested. Selectivity was determined based on the number of MEA channels that recorded a CAP in response to abdominal VNS through only one cuff contact pair. Further, the somatotopic organization of the responses in the nodose ganglia was determined, and it was demonstrated that low amplitude stimulation through different cuff contact pairs can evoke unique effects on gastric myoelectric activity.

**Figure 1.**
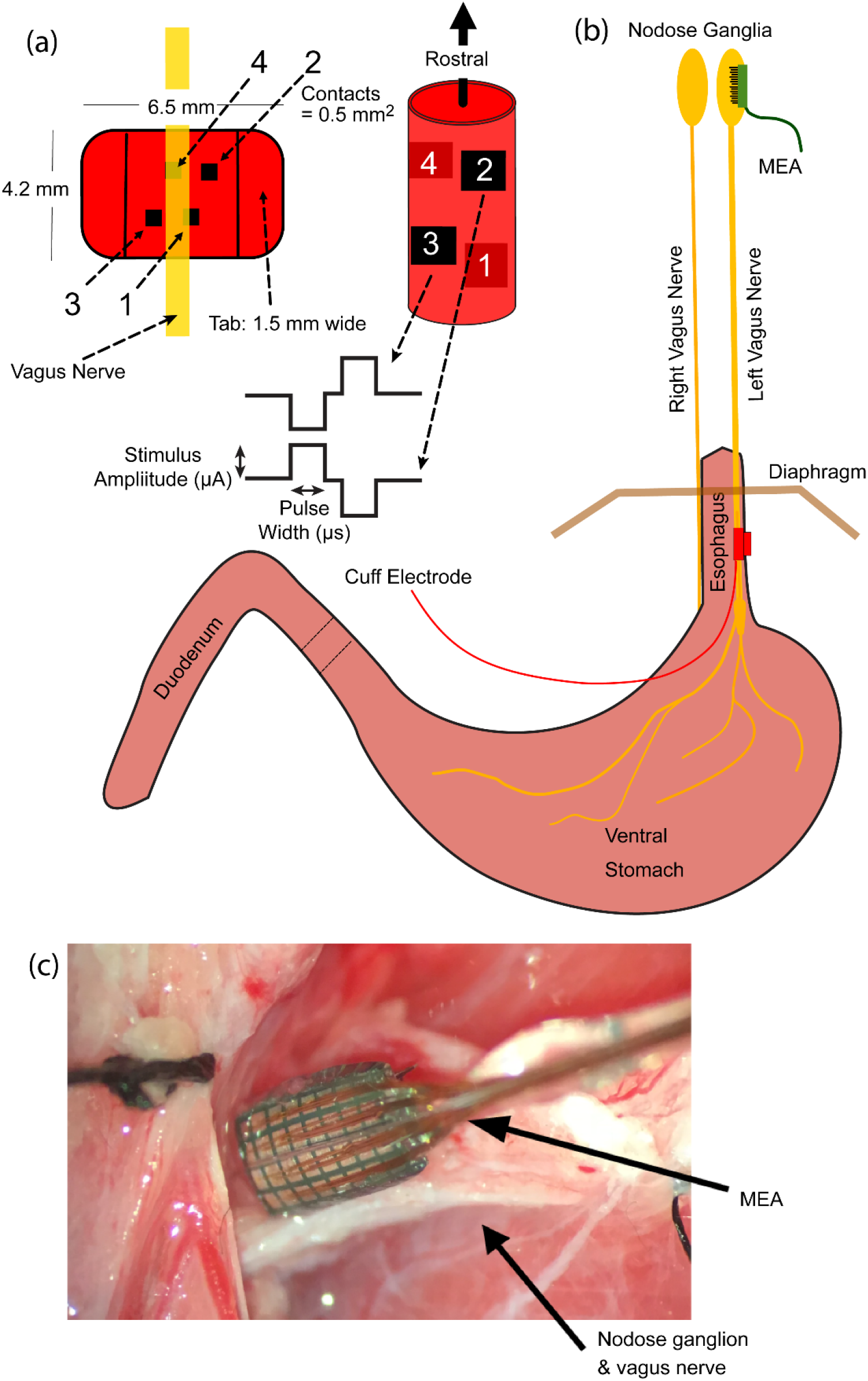
Experimental setup. In six ferrets, (a,b) a four-contact cuff electrode was wrapped around either the ventral or dorsal abdominal vagus nerve to deliver bipolar symmetric stimulus pulses. (b,c) A 32-channel microelectrode array (MEA) was inserted into the left or right nodose ganglion to record evoked responses to stimulation of the abdominal vagus.

**Figure 2.**
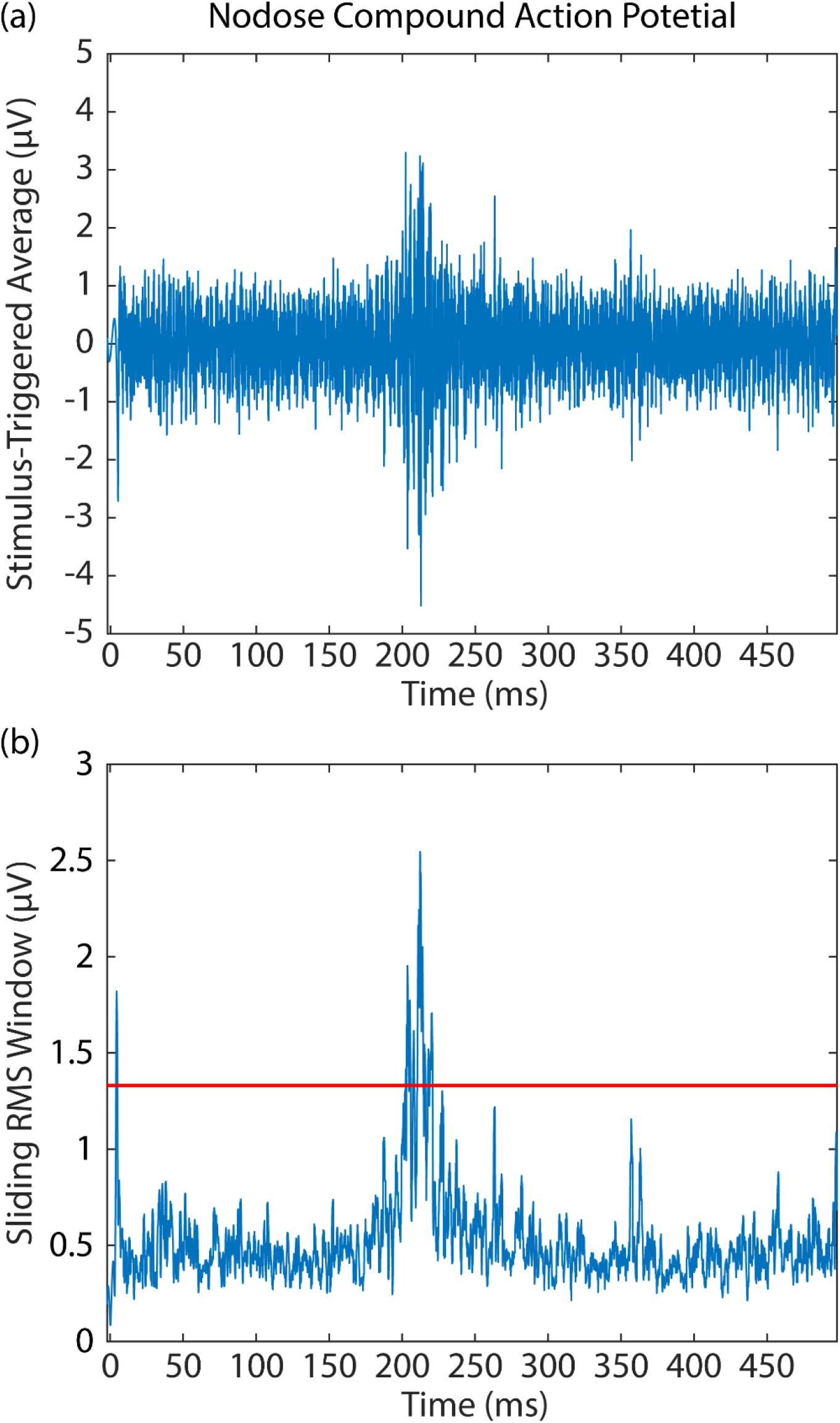
Nodose compound action potentials. (a) 120 repetitions of a single stimulus pulse were delivered at 2 Hz and a stimulus-triggered average was calculated to reduce noise amplitude and improve signal-to-noise ratio. (b) A sliding 1 ms RMS window smoothed the averaged signal to further reduce noise. Detection threshold (red line) was set to 2.4-2.6 times the standard deviation of the smoothed, averaged signal.

### Abdominal VNS evokes CAP responses in the nodose ganglion

In all six animals, abdominal VNS through each cuff pair evoked CAP responses in multiple MEA channels. For pulse widths of 0.1, 0.5, and 1.0 ms, Table 1 shows the threshold stimulation amplitude that evoked a response in at least one MEA channel. It is important to note that we did not control the rotational orientation of the cuff on the vagus nerve, so the designation of cuff contact pairs 1:2 and 3:4 is arbitrary. All animals exhibited responses to stimulation with a pulse width of 0.1 ms, except one cuff pair in animal F22-19 and both cuff pairs in animal F25-19. In both of those animals, stimulation with a pulse width of 0.4 ms evoked responses in multiple MEA channels. Mean thresholds were 1027, 307, and 248 μA for pulse widths of 0.1, 0.5, and 1.0 ms, respectively, in accordance with the expected strength-duration relationship in which threshold amplitude decreases exponentially as pulse width increases^25^.

**Table 1:** Threshold amplitude (μA) for each pulse width. NR: no response was detected at any amplitude, up to 3 mA for this pulse width. NT: stimulation at this pulse width was not tested in this animal.

|  | 0.1 ms |  | 0.4 ms |  | 0.5 ms |  | 1.0 ms |  | Vagus nerve trunk stimulated |
| --- | --- | --- | --- | --- | --- | --- | --- | --- | --- |
|  | Cuff contact pair |  |  |  |  |  |  |  |  |
| Animal | 1:2 | 3:4 | 1:2 | 3:4 | 1:2 | 3:4 | 1:2 | 3:4 |  |
| **F21-19** | 400 | 400 | NT | NT | 620 | 140 | 460 | 140 | Ventral |
| **F22-19** | 1500 | NR | NT | 1500 | 400 | 400 | 400 | 400 | Ventral |
| **F19-19** | 1500 | 1500 | NT | NT | 400 | 400 | 220 | 220 | Ventral |
| **F25-19** | NR | NR | 320 | 180 | 140 | 140 | 140 | 140 | Dorsal |
| **F26-19** | 1500 | 1500 | NT | NT | 400 | 400 | 400 | 220 | Dorsal |
| **F34-19** | 540 | 400 | NT | NT | 140 | 100 | 140 | 100 | Dorsal |

### Selectivity of abdominal VNS

To quantify selectivity of abdominal VNS, we characterized recruitment within the MEA (Figure 3) as stimulation amplitude increased from threshold. We also calculated a selectivity index (SI), which, when maximized, selects stimulation parameters for each pair of cuff contacts that maximize the number of MEA channels with evoked responses while minimizing the number of overlapping MEA channels that respond to stimulation through both cuff contact pairs. Table II shows the stimulation amplitudes that maximized SI for each cuff contact pair at each pulse width in all animals. For all animals (Figure 4), as amplitude or pulse width increased, stimulation evoked responses on multiple additional MEA channels across the entire array. Maximizing the selectivity index while limiting overlap to no more than 3 MEA channels (i.e. 10% of 32 total channels) demonstrated that stimulation through pairs of cuff contacts could selectively activate an average of 4, 6, and 6 MEA channels with pulse widths of 0.1, 0.5, and 1 ms, respectively. It is important to note, that even with 1 ms pulses at 3 mA, stimulation through an individual electrode typically evoked responses in ~20 MEA channels (gray bars in Figure 4), with some MEA channels never detecting any CAP response to stimulation.

**Figure 3.**
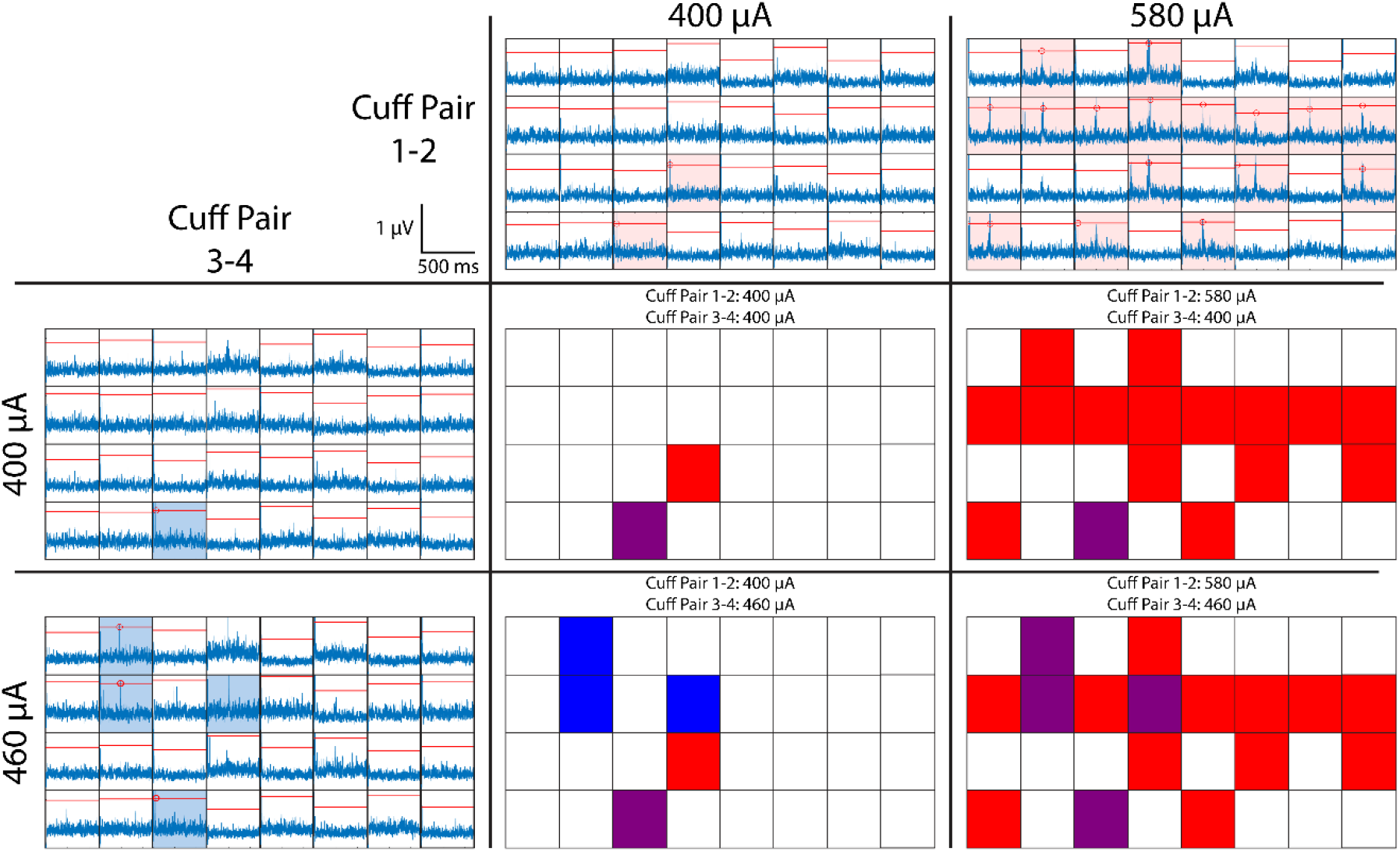
Selectivity as a function of stimulation amplitude. Grids of blue traces show 500 ms post-stimulation recordings from 32 MEA channels, with red horizontal lines representing the threshold for detecting an evoked response. Traces with red shading include a response evoked by stimulation from cuff pair 1-2. Traces with blue shading include a response evoked by stimulation from cuff pair 3-4. Grids of red, blue, and purple squares show the pattern of recruitment and overlap in responses at various stimulation amplitudes. Purple squares represent a MEA channel with a response evoked by stimulation from both cuff pairs. In this example from a single animal, as stimulation amplitude increases from 400 μA to 580 μA for bipolar cuff pair 1-2, the number of selectively responding MEA channels increases from one to fifteen. Similarly, as stimulation amplitude increases from 400 μA to 460 μA for bipolar cuff pair 3-4, the number of selectively responding MEA channels increases from one to three. Additionally, as stimulation amplitude increases, the number of MEA channels responding to both bipolar cuff pairs increases from one to four.

**Figure 4.**
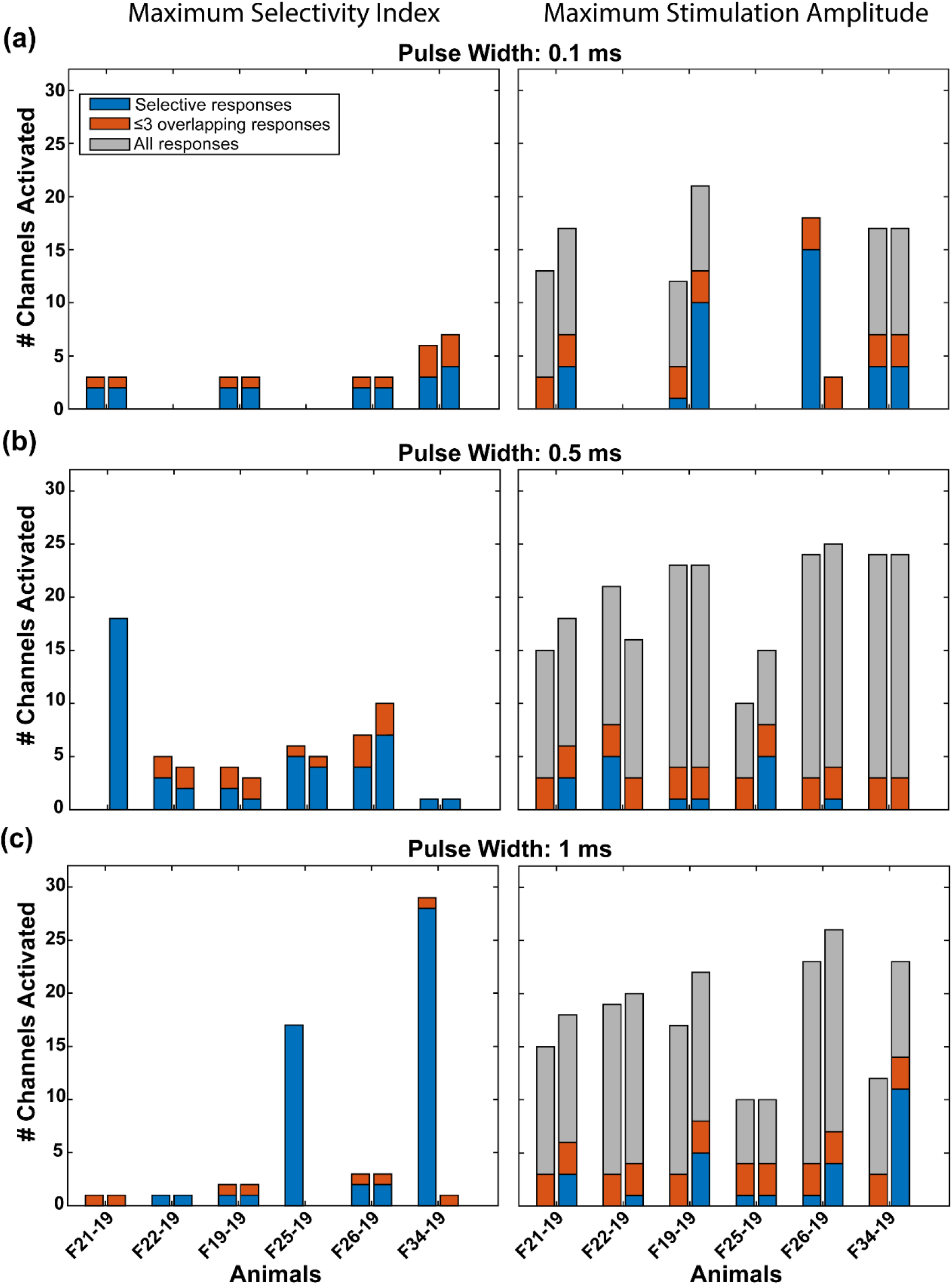
Selectivity of abdominal VNS. Number of responding MEA channels for each animal at (left) the stimulation amplitude that maximized SI and (right) the maximum tested stimulation amplitude (i.e. 3 mA). Pairs of bars represent the two bipolar pairs of cuff contacts in each animal (left is cuff pair 1-2, right is cuff pair 3-4). Blue bars represent MEA channels responding to only one of the two bipolar pairs, orange bars allow for up to 3 overlapping channels (i.e. 10% overlap), and gray bars include all non-selectively stimulated channels. Stimulation pulse widths include (a) 0.1 ms, (b) 0.5 ms, and (c) 1 ms.

**Table 2:** Stimulation amplitudes (μA) that maximized selectivity index for each pulse width. NR: no response was detected at any amplitude, up to 3 mA for this pulse width for at least one bipolar contact pair. NT: stimulation at this pulse width was not tested in this animal.

| Animal | 0.1 ms |  | 0.4 ms |  | 0.5 ms |  | 1.0 ms |  |
| --- | --- | --- | --- | --- | --- | --- | --- | --- |
|  | Cuff contact pair |  |  |  |  |  |  |  |
|  | 1:2 | 3:4 | 1:2 | 3:4 | 1:2 | 3:4 | 1:2 | 3:4 |
| F21-19 | 540 | 460 | NT | NT | 680 | 400 | 760 | 1500 |
| F22-19 | NR | NR | NT | NT | 580 | 3000 | 1500 | 760 |
| F19-19 | 2080 | 2080 | NT | NT | 500 | 640 | 400 | 380 |
| F25-19 | NR | NR | 380 | 180 | 220 | 140 | 140 | 140 |
| F26-19 | 1500 | 3000 | NT | NT | 760 | 580 | 500 | 280 |
| F34-19 | 3000 | 640 | NT | NT | 140 | 100 | 3000 | 140 |

### Conduction velocities of VNS evoked responses

To determine the types of nerve fibers (e.g. Aδ vs C) activated by abdominal VNS, the conduction velocities of the evoked CAPs in the nodose ganglion were calculated (Figure 5). At both threshold and the maximum stimulation amplitude tested (i.e. 3 mA), the majority of evoked responses (70.1% and 91.7%, respectively) had conduction velocities between 0 and 3 m/s (i.e. C fibers)^26^. At the maximum tested stimulation amplitude, other groups of neurons were activated with conduction velocities of 6-10 m/s and 12-14 m/s. While we did not quantify the receptive fields or modalities of these neurons, their conduction velocities suggest C-fibers were primarily recruited, along with a smaller population of Aδ fibers.

**Figure 5.**
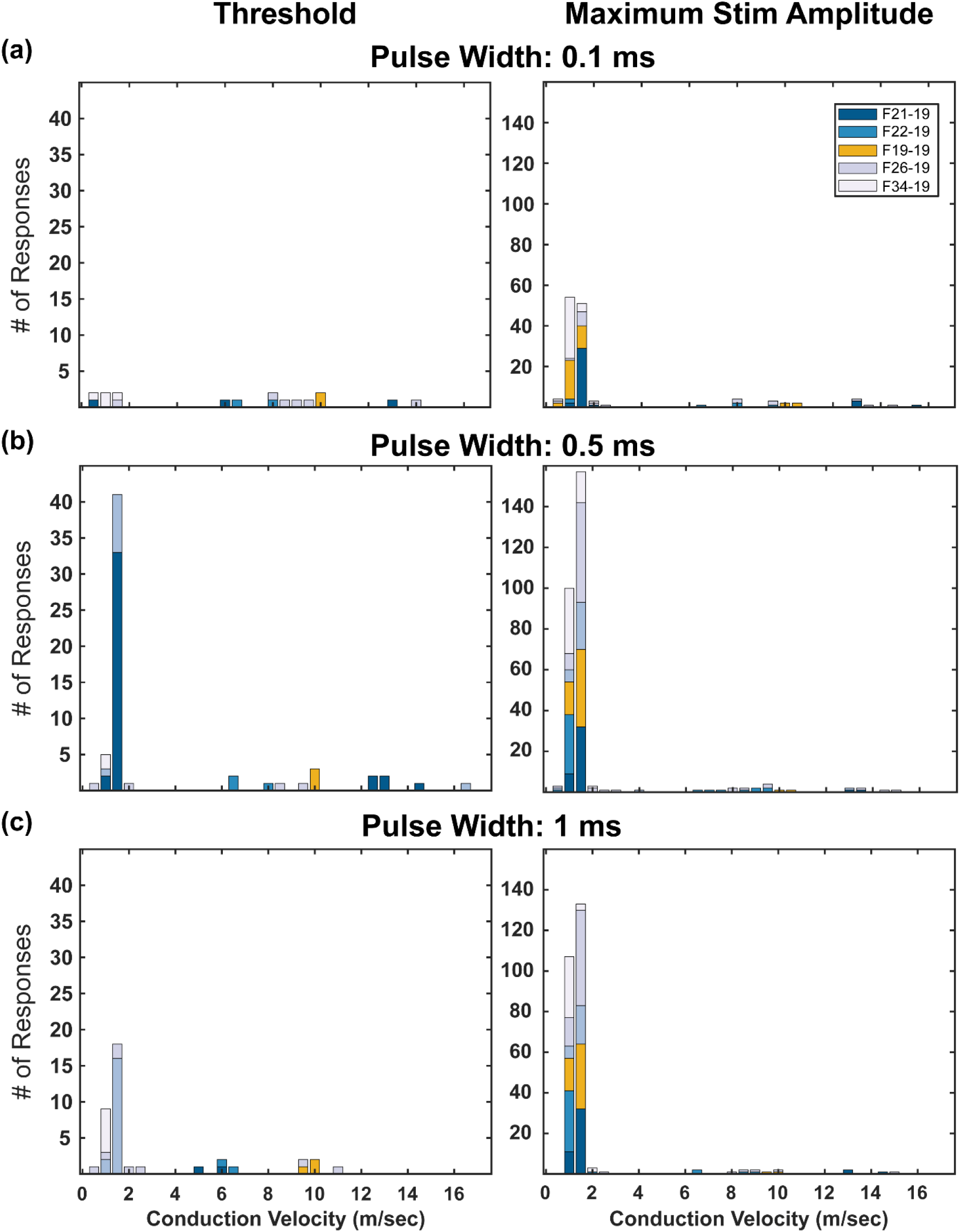
Conduction velocities of evoked responses. Histograms of the conduction velocities of evoked responses at (left) threshold and (right) maximum stimulation amplitude (i.e. 3 mA) for pulse widths of 0.1, 0.5, and 1 ms. Most evoked responses had conduction velocities between 0 and 3 m/s (i.e. c fibers), although a smaller set of responses had conduction velocities between 3 and 30 m/s (i.e. Aδ fibers).

### Patterns of recruitment across the nodose ganglion

To understand if there is somatotopic organization of the nodose ganglion with respect to the abdominal vagus nerve, we performed a nearest neighbor analysis for the MEA channels that responded to stimulation for each animal. Trials where at least one of the cuff pairs elicited no selective responses at the stimulation amplitude that maximized SI were excluded from this analysis. MEA channels that were activated by both cuff pairs were also excluded from this analysis. Histograms of this analysis across 6 ferrets are shown in Figure 6A.

**Figure 6.**
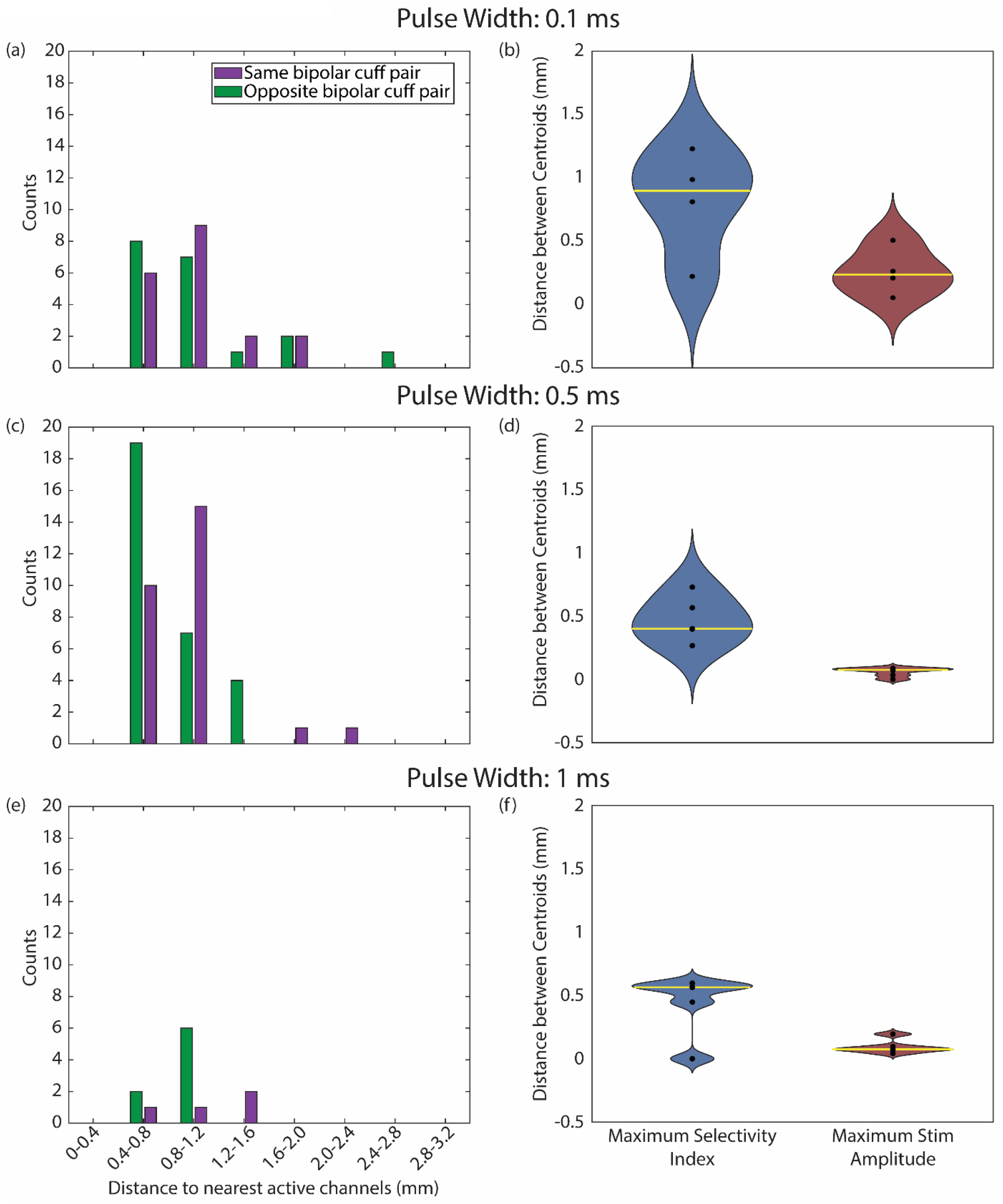
Distance between responding MEA channels. (a,c,e) Distance between each responding MEA channel and the nearest responding MEA channel from the same bipolar cuff pair (purple) or the other bipolar cuff pair (green) for pulse widths of (a) 0.1, (c) 0.5, and (e) 1 ms at the stimulation amplitude that maximized SI. (b,d,f) Distance between centroids of activation on the MEA for stimulation through each of the bipolar cuff pairs at the stimulation amplitude that maximized SI (blue) and the maximum tested stimulation amplitude (i.e. 3 mA; red) for pulse widths of (b) 0.1, (d) 0.5, and (f) 1 ms. Yellow lines show the median distance.

To further understand how recruitment of neurons at the nodose ganglion changed as a function of stimulation amplitude, we calculated the centroid location off all responding MEA channels for each bipolar cuff pair. We calculated these centroids at the stimulation amplitude that maximized SI and at the maximum stimulation amplitude tested (i.e., 3 mA). We then calculated the distance between the centroid of each cuff pair and compared these distances to determine if there was a change in the overall pattern of activation across the MEA as a function of stimulation amplitude (Figure 6B). For all animals and pulse widths, a Wilcoxon signed rank test shows that there was no statistical difference between conditions (p > 0.05 for all tests).

### Functional effects of selective abdominal VNS

In a separate set of experiments in three additional ferrets, we examined the effects of abdominal VNS on GI myoelectric activity. The goal of these experiments was to determine if low amplitude VNS through bipolar pairs of cuff contacts on the abdominal vagus nerve could selectively drive functional changes in the stomach. For these experiments, a four-contact cuff was implanted around the abdominal vagus and four four-contact paddle electrodes were sutured to the serosal surface of the stomach to record GI myoelectric activity. Across the three ferrets, 10 GI myoelectric signals displayed a statistically significant dominant frequency peak at baseline (9.62 ± 0.63 counts per minute; cpm). In all ferrets, VNS through at least one of the bipolar cuff pairs selectively evoked a distinct effect on normal GI myoelectric activity. VNS was delivered at a frequency of 15 Hz and a pulse width of 0.1 ms in all ferrets. For Ferret 15-18, when stimulation amplitude was 0.1 mA, there was no change in the GI myoelectric activity for either bipolar pair. At 0.2 mA, stimulation on one pair of bipolar contacts (1-2) did not result in any change in the GI myoelectric activity, while stimulation on the other pair of bipolar contacts (3-4) resulted in a decrease in the power in the normogastric range (8-11 cpm) to nearly 0 (Figure 7A). This change in GI myoelectric activity was not accompanied by any overt behavioral response (retching or emesis). The waterfall plots in Figure 7B display the change in the power spectral density of the signal in response to stimulation on bipolar contacts 1-2 and 3-4. For stimulation on contacts 3-4 we observed an immediate suppression of GI myoelectric activity upon stimulation onset.

**Figure 7.**
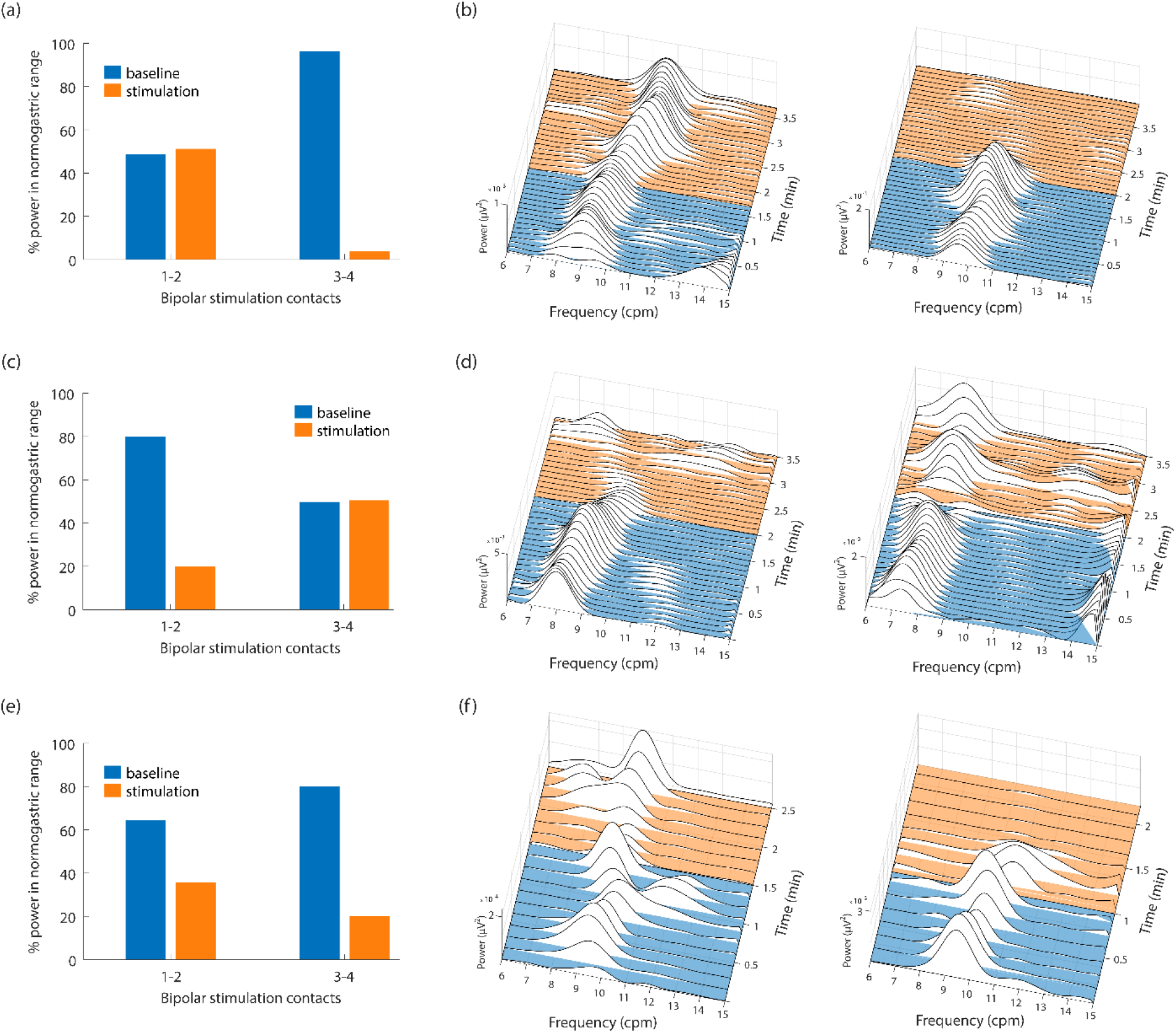
Low amplitude abdominal VNS selectively changes GI myoelectric activity. (a,c,e) In three animals, stimulation (orange) through one cuff pair caused a substantial change in normogastric power as compared to baseline (blue), while stimulation through the other cuff pair had a smaller or no effect. (b,d,e) Waterfall plots show signal power in frequencies between 6 and 15 CPM over time. Blue lines are pre-stimulation baseline and orange lines are during stimulation.

For Ferret 13-18, there was no change in the GI myoelectric activity for either bipolar pair when the stimulation amplitude was below 1 mA. At 1 mA, stimulation on one pair of bipolar contacts (1-2) resulted in a decrease in the power in the normogastric range (8-11 cpm) to 20% of the total power in the recorded signal (Figure 7C). Approximately 1.5 minutes after the onset of stimulation we observed retching followed by emesis. The waterfall plot in Figure 7D displays the change in the power spectral density of the signal leading up to the first retch, in response to stimulation on bipolar contacts 1-2. Stimulation on the other pair of bipolar contacts (3-4) did not result in any change in the GI myoelectric activity and was not accompanied by any overt behavioral response.

Similarly, for Ferret 14-18, there was no change in the GI myoelectric activity for either bipolar pair when the stimulation amplitude was below 1 mA. At 1 mA, stimulation on one pair of bipolar contacts (3-4) resulted in a decrease in the power in the normogastric range (8-11 cpm) to 20% of the total power in the recorded signal (Figure 7E). This decrease in power was accompanied by retching and emesis approximately 1.1 minutes after the onset of stimulation. The waterfall plot in Figure 7F displays the change in the power spectral density of the signal leading up to the first retch, in response to stimulation on bipolar contacts 3-4. Stimulation at 1 mA on the other pair of bipolar contacts (1-2) resulted in a decrease in the power in the normogastric range (8-11 cpm) to 38% of the total power in the recorded signal. However, this was not accompanied by any overt behavioral response (retching or emesis). For ferrets 15-18 and 13-18, the decrease in power in the normogastric range, roughly coincided with the onset of stimulation whereas for Ferret 14-18, the suppression of GI myoelectric activity occurred 20-30 seconds after the onset of stimulation.

## Discussion

The goal of this study was to selectively stimulate sub-populations of fibers within the abdominal vagus nerve to drive changes in GI activity that could be used for treating diseases such as gastroparesis and obesity. In six ferrets, we demonstrated that a multi-contact cuff electrode wrapped around the abdominal vagus nerve can selectively activate independent populations of nerve fibers, and in three other ferrets, we showed that selective VNS can drive changes in GI myoelectric activity. We assessed the selectivity of stimulation by recording evoked CAPs through a MEA inserted into the nodose ganglion of the vagus and quantifed the number of microelectrode channels activated by one or both bipolar pairs of cuff contacts. In all animals except one (F21-19), threshold-level stimulation through both cuff pairs drove selective activation of fibers recorded by at least one MEA channel. Pulse width had a critical effect on the ability to selectively activate axons in the abdominal vagus nerve. At a short pulse width (i.e. 0.1 ms), evoked CAPs were recorded from relatively few MEA channels, with no responses in two animals at maximal stimulation amplitude. Conversely, at a long pulse width (i.e. 1 ms), evoked CAPs were recorded from most MEA channels, although very few were selectively activated by only one of the bipolar cuff pairs. A pulse width of 0.5 ms produced a balance between these two extremes and the most selective activation of MEA channels, with 1-7 MEA channels selectively activated by each bipolar cuff pair in each animal.

Abdominal VNS primarily activated fibers with conduction velocities below 3 m/s, although there were also responses from a smaller set of fibers with faster conduction velocities. The abdominal vagus nerve contains both afferent and efferent pathways that convey a variety of information to and from the abdominal organs^27–29^. Previous studies have shown that the nerve is comprised primarily of Aδ and C fibers, which have conduction velocities of 3-30 and 0-3 m/s, respectively^26^. The larger diameter Aδ fibers are expected to have a lower threshold for extracellular electrical stimulation, although the much higher prevalence of C fibers was likely the primary reason we observed so many more responses from this fiber type^27,30^.

We also examined the patterns of activation across the MEA to determine if there was a structural relationship between the location of stimulation at the abdominal vagus nerve and the location of activation in the nodose ganglion. If there was a strong somatotopic relationship between the abdominal vagus nerve and the nodose ganglion, we would expect that the MEA channels with CAPs evoked by one bipolar stimulation pair would be spatially separate from those channels with CAPs evoked by the other bipolar stimulation pair. Instead, we found that there was no difference between the distances of MEA channels with responses evoked by the same bipolar pair and those evoked by different bipolar pairs. Further, high amplitude stimulation at the abdominal vagus nerve evoked responses on channels throughout the MEA, and there was no statistical difference in the location of centroids of MEA channels recording responses from the two bipolar cuff pairs, either for stimulation that maximized SI or at the maximum tested amplitude. These results suggest that it may be challenging to achieve selective recruitment of abdominal vagus nerve fibers with stimulation at the nodose ganglion because there is not substantial somatotopic organization within that structure.

In addition to quantifying the selectivity of abdominal VNS, we also measured its effect on GI function through changes in GI myoelectric activity. In three animals, we demonstrated that stimulation through one bipolar pair of cuff contacts had little or no effect on myoelectric activity, while stimulation through the other pair of contacts strongly suppressed that activity. These preliminary results support that VNS can have selective and differential effects on the GI system. It is unclear whether the effects of abdominal VNS on gastric myoelectric responses is due to stimulation of afferent or efferent vagal pathways; indeed, the vagus nerve contains vago-vagal reflex pathways that project to the hindbrain and return to the periphery^31^.

While these results demonstrate the promise of abdominal VNS for selectively stimulating the GI system, several important limitations should be addressed. Even when we maximized SI, selective responses were still limited to 1-7 MEA channels per bipolar cuff pair (i.e. ^~^3-20% of all MEA channels). Still, this level of selective activation may be sufficient to achieve functional effects, as was evident in three animals in which stimulation selectively evoked a substantial change in GI myoelectric activity. Multiple factors may have limited the selectivity of abdominal VNS in these experiments. The binary search method used here is an efficient approach for quickly finding threshold, but it results in irregular sampling of the parameter space that likely limited our ability to optimize stimulation parameters *post hoc*. Future work should use parameter space sampling methods that are specifically designed for optimizing stimulation parameters for selectivity^9^. Additionally, while the ferret is a good model for GI function (e.g. because of the intact emetic response), the ferret abdominal vagus is small (500-750 μm diameter) and monofascicular, which likely limited our ability to achieve selective stimulation. Future work should focus on animal models with vagus nerve anatomy similar to humans, such as the pig^32^.

In addition to issues of VNS selectivity, there are broader limitations of the current study, including effects of anesthesia and physiological variability across animals. Our studies were performed using inhalational isoflurane. Isoflurane is known to dampen activity of neuronal cell bodies in peripheral ganglia^33^, which could have biased our data. Although this is likely not a problem in the context of VNS trials because even quiescent neurons can be stimulated, we did use basal neural activity to determine if the MEA insertion into the nodose ganglia had adequate numbers of active channels for VNS testing. Moreover, it is not well documented but isoflurane is reported to affect gastric myoelectric activity^34^. There are also substantial physiological differences between animals, which we have observed in in gastric myoelectric activity in awake and anesthetized ferrets^35^. In this study, we quantified selectivity via CAP recordings in one set of ferrets and measured changes in GI myoelectric activity with similar stimulation parameters in a different set of animals. This approach was necessary because implantation of the MEA in the nodose may have disrupted GI myoelectric activity, but it required us to quantify selectivity and measure functional changes in separate animals. Future studies should demonstrate selectivity, optimize VNS, and quantify functional effects in each subject.

Our focus in the current study, and the clinical impact, is the optimization of VNS selectivity. Current commercial cuff electrode geometry is limited in the number of circumferential contacts, and thus selectivity. In the ferret, we have tested 4, 6, and 8 contact vagus nerve cuff electrodes, and 6 contacts (two rows of 3 circumferential contacts) appear to be the current limit for consistent manufacturing quality for abdominal vagus nerve of the ferret. This limitation further justifies that future experiments should be conducted in species with larger vagus nerve diameters, such as the cat or pig. In addition, there is potential to increase the efficiency of algorithmic optimization of VNS. We used recordings of the nodose ganglia to assess selectivity but it is unlikely this approach could be applied clinically; therefore, it will be important to use more accessible targets, such as serosal gastric myoelectric activity or cuff electrode recordings from the vagus nerve to confirm VNS selectivity [e.g., 3, 4]. Such an approach may lead to the application of closed-loop approaches to record and apply VNS to treat diseases of the abdominal cavity, such as GI disease, obesity, diabetes, and inflammatory disorders.

## Methods

### Animals

Nine adult male ferrets (weight: 1-1.7 kg; Marshall BioResources, North Rose, NY, USA) were used in this study. All experimental procedures were approved by the University of Pittsburgh Institutional Animal Care and Use Committee. Animals were housed in wire cages (62 x 74 x 46 cm) under a 12-hour standard light cycle (lights on at 0700 h), in a temperature (20-24°C) and humidity (30-70%) controlled environment. Food (ferret kibble: Mazuri Exotic Animal Nutrition, St. Louis, MI) and drinking water were freely available. Food was removed 3 hours before induction of anesthesia. After each experiment, animals were euthanized with an injection of a 5 ml solution of SomnaSol (390 mg/ml pentobarbital sodium; 5 mg/ml phenytoin sodium; SomnaSol EUTHANASIA-III Solution, Henry Schein Animal Health, Dublin, Ohio, USA).

### Surgical procedure

Anesthesia was induced and maintained with inhaled isoflurane (5% induction, 1-3% maintenance), and a tracheotomy was performed followed by insertion of an intratracheal tube to monitor respiration and deliver the anesthetic agent. Vital signs were monitored throughout the experiment, including blood pressure, heart rate, body temperature, and respiration rate, and isoflurane was adjusted to maintain a surgical plane of anesthesia (i.e., non-responsive to toe pinch). In six of the ferrets, a laparotomy was performed, followed by implantation of a four-contact nerve-cuff electrode (Micro Leads, Inc. Somerville, MA) around the ventral (n=3) or the dorsal (n=3) abdominal vagus trunk (Figure 1). The cuff electrode contacts (area: 5 mm^2^, spaced 1 mm x 0.6 mm, 600 μm inner diameter) were arranged in two bipolar pairs with electrodes in each pair offset from each other by 90 degrees, and the pairs spaced equally around the circumference of the nerve to provide current steering for targeted stimulation. A 32-channel MEA (4-by-8 electrode arrangement with 400 μm inter-electrode pitch, 1 mm long shanks; Black-Rock Microsystems, Salt Lake City, UT) was implanted into the nodose ganglion with a pneumatic inserter for rapid insertion through the epineurium. Recordings of spontaneous single-unit activity were used to verify insertion of the MEA into the nodose, and if necessary, additional impacts were applied to insert the device further. A platinum wire was placed near the nodose to act as a reference and another platinum wire was inserted under the skin as the recording ground. In the additional three ferrets, an identical procedure was used to implant a four-contact nerve-cuff electrode around the ventral abdominal vagus nerve and four planar electrodes, each with four contacts (Micro Leads, Inc. Somerville, MA), were sutured to the ventral gastric surface and the duodenum. The locations these planar electrodes on the ventral gastric surface was similar to our prior study^35^.

### Stimulation and data acquisition

A Grapevine Neural Interface Processor (Ripple, Salt Lake City, UT) and stimulation headstage (Nano2+Stim) were used to deliver stimulation to the pairs of electrodes on the abdominal cuff, while a recording headstage (Nano2) was used to record evoked CAP signals from the nodose MEA and GI myoelectric activity from the serosal surface of the stomach. Nodose recordings were sampled at 30 kHz and filtered with a high pass filter at 150 Hz and low pass filter at 7500 Hz. GI myoelectric signals were sampled at 30 kHz with a high pass filter at 0.1 Hz and a low pass filter at 7500 Hz. An adapter was inserted between the cuff and stimulation headstage to connect four output channels in parallel to increase the maximum stimulation amplitude per channel from 1.5 mA to 6 mA. For all experiment sessions, the impedance of electrodes was measured at 1 kHz pre- and post-implantation as well as after each recording session to track changes in the electrode-tissue interface and ensure the connectivity of the system.

### Stimulus-triggered averaging of compound action potentials

MEA recordings were analyzed using an automated algorithm written in Matlab (version 2017a, Mathworks, Natick, MA) to detect stimulation-evoked CAPs and minimize the time required to determine the parameters for threshold (amplitude and pulse width) for each bipolar cuff pair. For a given pulse width and amplitude combination, 120 pulses were delivered at 2 Hz through a longitudinal bipolar pair of electrodes in the cuff. The data recorded from each electrode within the MEA was segmented into a window around each stimulation event with a time duration of 2 ms pre-stimulation and 498 ms post-stimulation. The ensemble average of these windows was calculated to improve the signal-to-noise ratio. A sliding 1 ms moving root mean squared (RMS; Figure 2) window with a step size of 0.1 ms was used to smooth this stimulus-triggered averaged (STA) signal. To remove artifacts in the signal caused by EMG activity or missing data packets, blanking was implemented over portions of the signal crossing 8 mV with a linearly interpolated signal between the start and end of the artifact periods. Baseline noise levels were measured for each MEA channel from stimulation recordings by first blanking a window centered at each stimulation event with a duration of 3 ms longer than the pulse width and secondly calculating the ensemble average of 120 randomly selected windows from this signal. The threshold for detecting a CAP response was set to between 2.4 and 2.6 standard deviations above the mean of the ensemble average of 120 segments of baseline recording. This multiplier of standard deviation was determined by creating a subset of ground truth data from expert evaluation of MEA recordings and generating ROC curves from a range of gain values, and selecting a multiplier that achieved a false positive rate of less than 10%. For each bipolar pair of cuff electrode contacts, stimulation threshold was defined for each pulse width as the minimum amplitude required to activate at least one MEA channel.

Threshold amplitude was determined for three different pulse widths (0.1, 0.5, and 1 ms). In one animal with a ventral cuff and one animal with a dorsal cuff, we also tested 400 μs pulses because no response was measured at 0.1 ms with amplitudes up to 3 mA.

The conduction velocity (CV) of nerve fibers responding to stimulation was calculated using the distance between the cuff and the electrode on the ventral or dorsal trunk, divided by the time between the stimulation event and when the RMS signal crossed the detection threshold. Signals were divided into time windows corresponding to 0.5 m/s increments, and only a single response could be detected within each time window.

### Optimization of selectivity stimulation parameters

In order to quantify our ability to selectively stimulate subpopulations of neurons in the abdominal vagus with a multi-contact cuff electrode, we performed a post-hoc optimization of stimulation parameters to maximize the number of nodose ganglion MEA channels with a CAP while minimizing the number of overlapping MEA channels with CAPs driven by both pairs of cuff electrodes. For a given pulse width, we varied pulse amplitude and calculated a selectivity index function of the form:

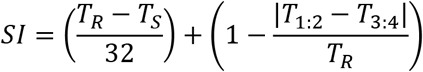

where SI is the selectivity index, T_R_ is the total number of nodose electrodes with an evoked CAP response to stimulation with either cuff contact pair, T_S_ is the total number of nodose electrode channels that recorded a response from stimulation through both cuff contact pairs (i.e., overlap in stimulation), T_1:2_ is the total number of nodose MEA channels with an evoked CAP response from stimulation through cuff contact pair 1:2, and T_3:4_ is the total number of nodose MEA electrodes that detected a response from stimulation through cuff contact pair 3-4. The first term in this equation calculates the number of MEA channels with non-overlapping responses to only one of the two cuff pairs, penalizing large overlaps in the number of MEA channels responding to stimulation through both cuff contact pairs. The second term penalizes imbalance between the responses from the two cuff contact pairs to prevent the condition in which stimulation through one cuff contact pair is maximized and no stimulation is delivered through the other cuff contact pair (i.e., it ensures that the responses are balanced between the two cuff contact pairs). An additional constraint was applied to severely penalize responses with more than three overlapping MEA channels (e.g. more than 10% overlap), to avoid producing results with excessive overlap that would not be functionally useful in tuning the effects of stimulation. By maximizing this SI equation, we determined the stimulation amplitude that maximized the number of selectively responding MEA channels.

### Nearest-neighbor analysis to quantify somatopic organization of nodose ganglion

To quantify the effect of stimulation amplitude on the location of activated MEA channels in the nodose ganglion, a nearest-neighbor analysis was performed for each pulse width at the stimulation amplitude that maximized SI and at the maximum amplitude that was tested (i.e. 3 mA). Animals where the maximum SI resulted in no selective responses for at least one cuff pair were excluded. For each MEA channel that recorded a response to abdominal VNS, we calculated the Euclidian distance to the nearest channel that recorded a response due to 1) stimulation from the same bipolar cuff pair with the same stimulation parameters and 2) from the opposing bipolar cuff pair with the stimulation parameters that elicited the maximum SI response. MEA channels that recorded non-selective responses from both cuff pairs were excluded because the distance to the nearest responding channel from the opposing bipolar cuff pair would always be zero but the distance to the nearest responding channel from the same bipolar cuff pair would always be greater than zero for these electrodes, resulting in a non-informative skew in the result.

To further quantify changes in the location of activation across the nodose ganglion in response to abdominal VNS, the centroid of all MEA channels recording responses from VNS were calculated for each cuff pair at the stimulation amplitude that maximized SI and at the maximum stimulation amplitude tested (i.e. 3 mA). The distance from one corner of the MEA was calculated for each channel, and the average distances of all channels that recorded a response from abdominal VNS was calculated to determine the centroid of activation for each set of stimulation parameters. The Euclidian distance between these centroids for each cuff pair was then calculated to quantify the relative distance across the array between locations of activation for the two bipolar cuff pairs.

### Electrogastrogram as a functional measure of selectivity

We have previously shown that GI myoelectric activity can be used to identify the physiological state (normal, distended, pre-retch) of the stomach^35^. For this study, we analyzed data from a subset of those experiments to demonstrate the selective effect of VNS on GI myoelectric activity. Stimulation was delivered at the abdominal vagus through each bipolar cuff pair at a rate of 15 Hz with either 0.1 or 0.5 ms/phase symmetric pulses. Stimulation amplitude was increased on consecutive trials (0.2, 0.4, 0.6, 0.8, 1, 5 mA) until it resulted in retching. For each trial, GI myoelectric activity was recorded at baseline for 5 minutes followed by 2 minutes of stimulation. Trials where retching or emesis occurred were not included in the analysis.

GI myoelectric recordings were analyzed post-hoc using MATLAB (Mathworks, Natick, MA). For every planar electrode, the waveform recorded on each of the four contacts was averaged to generate a single GI myoelectric waveform for that electrode. Analysis methods for GI myoelectric activity were adopted from our prior study^35^. Briefly, each planar-averaged GI myoelectric signal was filtered using a low-pass Butterworth filter with a 2.5 Hz (150 cpm, 4^th^ order) cutoff. The filtered signal was then downsampled to 10 Hz and a second low-pass Butterworth filter with a cut-off frequency of 0.3 Hz (18 cpm, 2^nd^ order) was applied. GI signals that displayed a statistically significant dominant frequency in the 0 to 15 counts per minute (cpm) range were retained for all analysis. Each GI myoelectric signal was partitioned into 60-second segments with a 54-second (90%) overlap between consecutive segments and the power spectrum for each segment was computed using the fast Fourier transform (*fft*, bin size: 0.1 cpm). Additionally, the fraction of power in the normogastric range (8-11 cpm) during baseline and stimulation was compared to identify differential effects of vagus stimulation.

## Data Availability

All study data will be made available through the NIH SPARC data portal (https://sparc.science). We will also include a github repository with code to generate all figures from the manuscript from those data.

## Acknowledgements

This work was supported by National Institutes of Health funding (Common fund SPARC Program award U18TR002205).

## Author contributions

LEF, CCH, JAS, and ACN designed the experiments. CCH and DMM conducted the surgery. BJY, DMM, SF, and MS maintained anesthesia and monitored physiology. JO, LW, and BM designed and manufactured the multi-contact cuff electrodes. JAS, DWB, and ACN performed data analysis. JAS, DWB, ACN, CCH and LEF drafted the manuscript. All authors read and supplied edits for the final manuscript.

